# Nucleoporin1 maintains male germ unit organization and transport in Arabidopsis pollen tubes, likely through shaping nuclear morphology

**DOI:** 10.64898/2026.06.04.730007

**Authors:** Raj K Thapa, Gang Tian, Binghui Shan, Xin Xie, Qing Lu, Jie Shu, Chen Chen, Shaomin Bian, Xuyan Li, Sangeeta Dhaubhadel, Susanne E Kohalmi, Steven J Rothstein, Yuhai Cui

## Abstract

The male germ unit (MGU) in Arabidopsis pollen is comprised of one vegetative nucleus (VN) and two sperm nuclei (SN). It is evolutionarily specialized to deliver immotile sperm nuclei to an ovule for fertilization. Despite some progress in research on MGU, its organization and transport remain only partially understood. Here, we identified Nucleoporin1/136 as a new player in the structural organization and positioning of MGU in pollen tubes. We and others have previously reported the reduced fertility of *nup1-1* plants; however, the mechanism remains unknown. In this work, we further examined the role of *NUP1* in fertility using two mutant alleles, *nup1-1* and *nup1-2^-/+^*. The reciprocal crosses between the *nup1* mutants and the Col-0 wild type indicate that the *nup1* mutant pollen is defective. To study the effect of a complete *NUP1* knockout on pollen, we generated a transgenic line that produces pollen with and without *NUP1* expression. This work led to the surprising discovery that the NUP1 protein is inherited from the pollen mother cell to the daughter cell during microgametophyte development. Subsequent *in vitro* experiments showed that NUP1 is required for pollen germination and pollen tube elongation. Further microscopic studies demonstrated that NUP1 is highly expressed in VN and essential for maintaining nuclear shape and size. We also demonstrated that NUP1 is required for proper MGU organization and transport, likely through maintaining VN morphology. Notably, our finding of nuclear morphology-mediated regulation of MGU may also explain the mechanistic details underlying the defective MGU movement in previously reported mutants such as *kaku4*, *wit*, and *wip,* which have abnormal nuclear morphology.

## Introduction

An Arabidopsis pollen grain is composed of a vegetative nucleus and two immotile sperm nuclei, which are together called the male germ unit (MGU) (McCue et al., 2011). Unlike animal sperm cells, pollen sperm cells lack the self-propelling ability to reach the egg for fertilization. Therefore, pollen grows a long tube towards the female reproductive tissue and positions the sperm nuclei at its tip (Hamamura et al., 2011). After receiving signals from the female counterparts, the pollen tube bursts, releasing two sperm nuclei into the ovary to ensure double fertilization (Dresselhaus et al., 2016). One sperm nucleus fertilizes the egg cell, and the other fertilizes the central cell, resulting in the formation of the embryo and endosperm, respectively (Dresselhaus et al., 2016; Hamamura et al., 2011). This whole process, from pollen germination to fertilization, requires strong cell-cell communication (Zhong et al., 2025).

The vegetative nucleus is connected to the sperm nuclei and migrates as a unit within the pollen tube (Mogensen, 1992). Generally, the vegetative nucleus (VN) leads the movement, while the sperm nuclei (SN) trail behind (Zhou & Meier, 2014). Based on Arabidopsis T-DNA mutant studies, only a few proteins have been implicated in organizing MGU and in its migration to the pollen tube tip. Some forms of MGU organization and migration defects have been observed in pollen grain and pollen tubes of *wit12*, *wip123, kaku4*, *hug1, hug2,* and *callose synthase 3* mutants (Goto et al., 2020; Motomura et al., 2021; Yan et al., 2025; Zhou & Meier, 2014). The SN in the *wit12*, *wip123*, and *kaku4* mutants precede VN during their migration from the pollen grain to the tip of the pollen tube (Goto et al., 2020; Zhou & Meier, 2014). Identifying new players is obviously important for a full understanding of the MGU organization and its migration.

Pollen is the most vulnerable plant organ to heat stress (Chaturvedi et al., 2021). The MGU organization and transport can be disrupted by high temperatures, leading to fertilization failure and subsequent seed set impairment (Ge et al., 2011; Li et al., 2024). The average global temperature is set to rise 2-4 °C by the end of the 21^st^ century. Rising temperatures and frequent heat waves are likely to reduce the quality of pollens from crop plants, vegetables, and fruit trees. Therefore, a comprehensive understanding of the organization and transport of MGU in pollen and pollen tubes is critical for breeding heat-resistant crops.

Here, we show a novel function of Nucleoporin1 (NUP1)/NUP136 in the organization and transportation of MGU in pollen tubes. NUP1/NUP136 is a member of the Nuclear Pore Complex (NPC) that is composed of more than 30 types of proteins, and is located on the inner side of the nuclear envelope (Lu et al., 2010). NUP1 was also named NUP136 based on its molecular weight (Tamura et al., 2010). The canonical function of NPCs is the nucleo-cytoplasmic transport of mRNA and proteins (Fernandez-Martinez et al., 2017). However, many nucleoporins have been found to play multiple roles in plant growth and development, such as gene regulation, immune signalling, and plant fertility (Braud et al., 2012; Gu, 2018; Park et al., 2014). Of note, NUP1 was previously reported to have diverse functions, including mRNA export, regulation of cell size and nuclear morphology, and early seedling establishment (Lu et al., 2010; Tamura et al., 2010; Thapa et al., 2022; Thapa et al., 2026).

The *NUP1* gene is highly expressed in pollen (Winter et al., 2007), and its T-DNA knockdown allele, *nup1-1*, exhibits reduced fertility (Lu et al., 2010; Tamura et al., 2010). However, the heterozygous *nup1-2* ^-/+^ plant does not have any fertility defect (Lu et al., 2010). The *nup1-1* allele can be considered a weak allele because it still expresses some *NUP1* transcript (Tamura et al., 2010), while *nup1-2* is considered a strong allele because it is homozygous-gametophytic lethal and thus can only be maintained as heterozygous (Lu et al., 2010). Here, we used both T-DNA mutant lines, *nup1-1* and *nup1-2* ^-/+^ (Supplementary Figure 1), to study the role of NUP1 in male fertility. Our genetic, transgenic, cell-biological, and microscopic work revealed novel functions of NUP1 in the organization and transport of MGU in pollen tubes.

## Results

### 1. Reduced fertility in the *nup1* plants is due to defective male gametes

To study the role of NUP1 in fertility, the two mutant plants (*nup1-1* and *nup1-2* ^-/+^) were grown together with the Col-0 wild-type (Figure 1A). The *nup1-1* plants have shorter siliques and fewer seeds per silique compared to Col-0 (Figure 1B-D). The *nup1-2* ^-/+^ plants are heterozygous for a T-DNA insertion and appear similar to Col-0 in plant morphology and fertility (Figure 1A-D). The Col-0 and *nup1-2* ^-/+^ plants have a full seed set, but the *nup1-1* siliques contain some unfertilized ovules (Figure 1E and Supplementary Figure 2).

**Figure 1.**
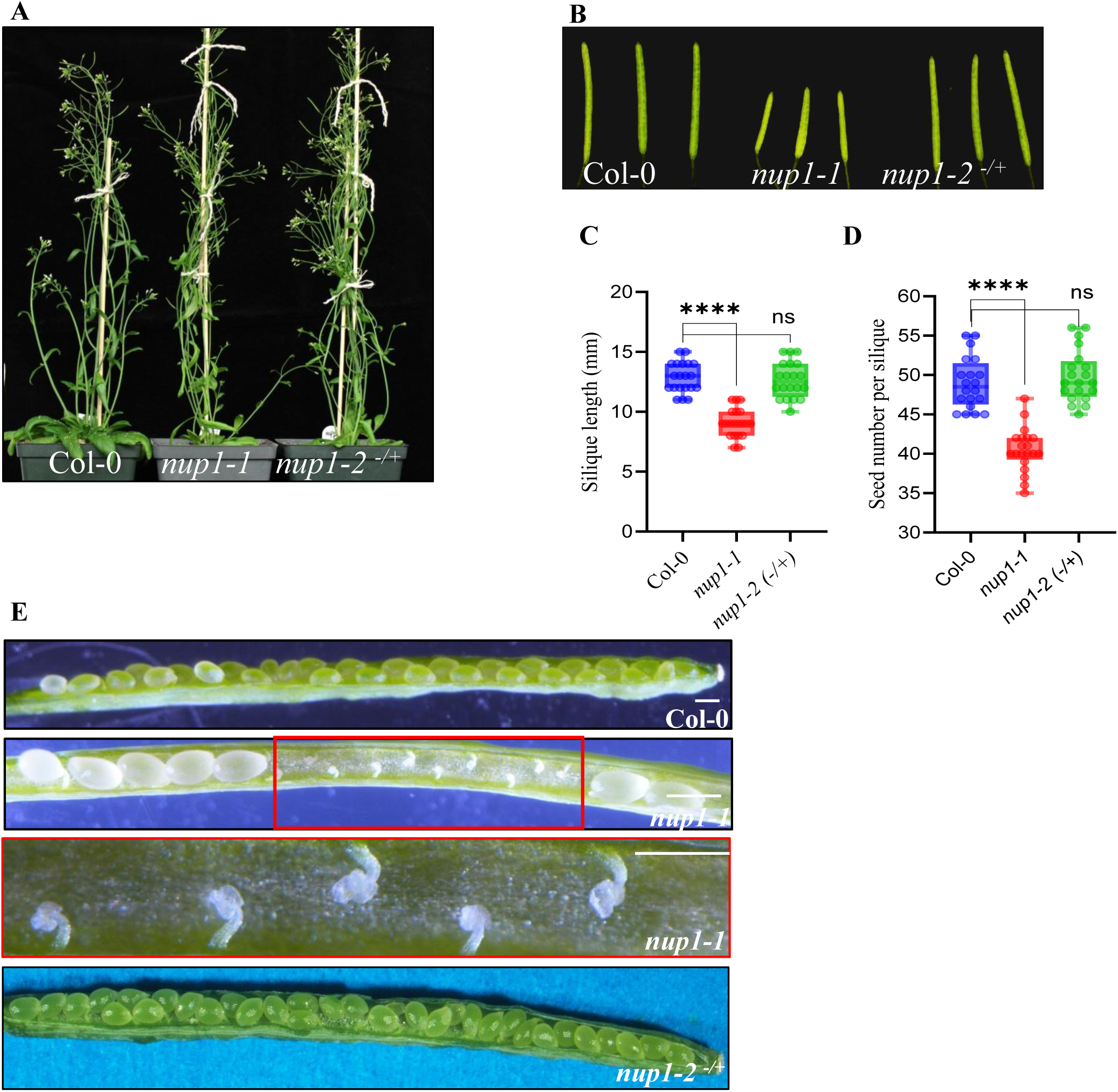
Comparison of fertility between Col-0, *nup1-1* and *nup1-2* (-/+) plants. **A.** Mature plant Col-0, *nup1-1* and *nup1-2* (-/+) **B.** Comparison of silique from Col-0, *nup1-1* and *nup1-2* (-/+) plants **C-D**. Silique length (C) and seeds per silique (D) of Col-0, *nup1-1* and *nup1-2* (-/+) **E.** Dissection of siliques for viewing seeds or unfertilized ovules.

To determine any fertility defects in pollen or ovules due to the knockout (ko) of *NUP1*, we conducted a reciprocal cross between Col-0 and *nup1-2* ^-/+^ plants. When Col-0 was used as a pollen donor, and *nup1-2* ^-/+^ as a recipient, the F2 seedlings were segregated as 1:1 (Wild type: *nup1-2* ^-/+^) (Supplementary Table 1). However, when reciprocal crosses were conducted, none of the progeny seedlings tested carried the *nup1-2* mutation. Also, self-crossing of *nup1-2* ^-/+^ resulted in a 1:1 ratio of wild type: *nup1-2* ^-/+^ (Supplementary Table 3). These genetic crosses suggested that the transmission of *nup1-2* ko pollen is completely blocked, while the *nup1-2* ko ovule is functionally unaffected.

### 2. The NUP1 protein is inherited from the pollen mother cell to the daughter cell during sporogenesis

The *nup1-2* ^-/+^ heterozygous plant is expected to produce pollen with and without *NUP1* expression based on the Mendelian segregation. However, there was no way to distinguish wild type from mutant pollen to study the effect of *NUP1* knockout on pollen. Therefore, we utilized the transgenic plant *nup1-2 ^-/-^ pNUP1:NUP1-YFP*, which is homozygous for both the NUP1-YFP transgene and the *nup1-2* mutation (hereafter referred to as NUP1-YFP (+/+)) (Lu et al., 2010). This plant, which has fertility similar to that of the Col-0 wild type (Supplementary Figure 3), was crossed with a *nup1-2 ^-/+^* heterozygous plant. Then, progenies were selected that are homozygous for the *nup1-2* T-DNA insertion by PCR-based genotyping and heterozygous for transgene (NUP1-YFP) expression by observing YFP-fluorescent/non-fluorescent pollen under a microscope. Such plants (hereafter referred to as NUP1-YFP (+/-)) produce two types of pollen: 1) NUP1-YFP (+), which is NUP1 complemented, and 2) NUP1-YFP (-), which is essentially *nup1-2*, in equal numbers in theory (Figure 2A). This allows for a side-by-side visual comparison of any developmental defects resulting from the *nup1-2* mutation. The transgenic plant, NUP1 YFP (+/+), which produces all wild-type-equivalent pollen with fluorescent NUP1-YFP (+), was used as a control (Supplementary Figure 4A-B). In the NUP-YFP (+/-) plants, we hypothesized that one Pollen Mother Cell (PMC) would produce two YFP-fluorescent (wild-type equivalent) and two non-fluorescent microspores (*nup1-2* mutant) in one tetrad (Figure 2A). Surprisingly, we observed that all four microspores in the tetrad showed YFP fluorescence, although the intensity was lower in two of them (Figure 2B, Supplementary Figure 5). However, only ∼50% of the mature pollen exhibited YFP fluorescence (Figure 2B). Upon further investigation, we found that NUP1 is expressed in almost all the unicellular stage pollen, but the NUP-YFP signal fades away in about ∼38% of the pollen at the bi-cellular stage (Figure 2C-D). By the time pollen matures (tricellular), ∼53% of them lose the fluorescence (Figure 2E-F). These results suggest that the NUP1 protein is inherited and gradually diluted during male gametophyte development.

**Figure 2.**
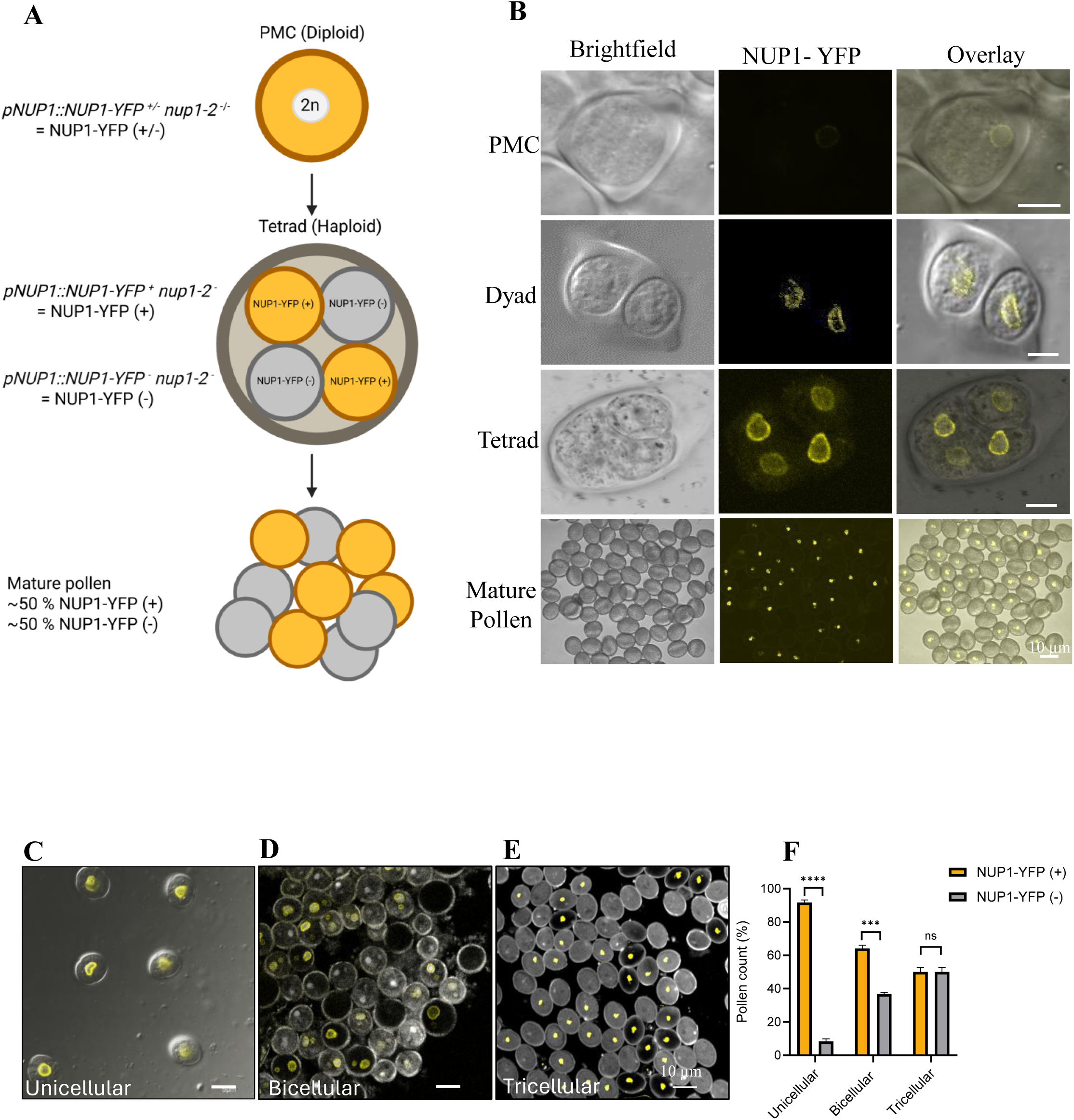
The NUP1 protein is inherited during male gametogenesis, and it partially compensates for *NUP1* genetic deficiency in pollen A. Schematic diagram of male gametogenesis in an Arabidopsis transgenic plant NUP1-YFP (+/-) producing ∼50% NUP1-YFP (+), and ∼50% NUP1-YFP (-) pollen. **B.** NUP1 expression pattern in NUP1-YFP (+/-) plants from pollen mother cell, dyad, tetrad and mature pollen stages. NUP1-YFP was expected to be observed in only half of the dyad or tetrad. Scale bar 10µm. **C-E.** NUP1 expression pattern in unicellular, bicellular and tricellular pollen stages of NUP1-YFP (+/-) plants. Scale bar 10µm. **F.** Proportion of NUP1-YFP (+) and NUP1-YFP (-) mature pollen in NUP1-YFP (+/-) plants. N= 200

### 3. NUP1 is required for pollen germination and elongation of the pollen tubes

Next, we conducted an *in vitro* pollen germination (PG) using pollen from NUP1-YFP (+/-) plants, which produce ∼50% NUP1-YFP (+) and ∼50% NUP1-YFP (-) pollen (Figure 3A-D). The NUP1-YFP (-) pollen has only half the germination rate of NUP1-YFP (+) (Figure 3D). Also, the NUP1-YFP (-) pollen has shorter pollen tube length (PTL) compared to that of NUP1-YFP (+) pollen (Figure 3E-F). Almost 30% of the NUP1-YFP (-) pollen seems to be arrested after the early phase of pollen tube elongation.

**Figure 3.**
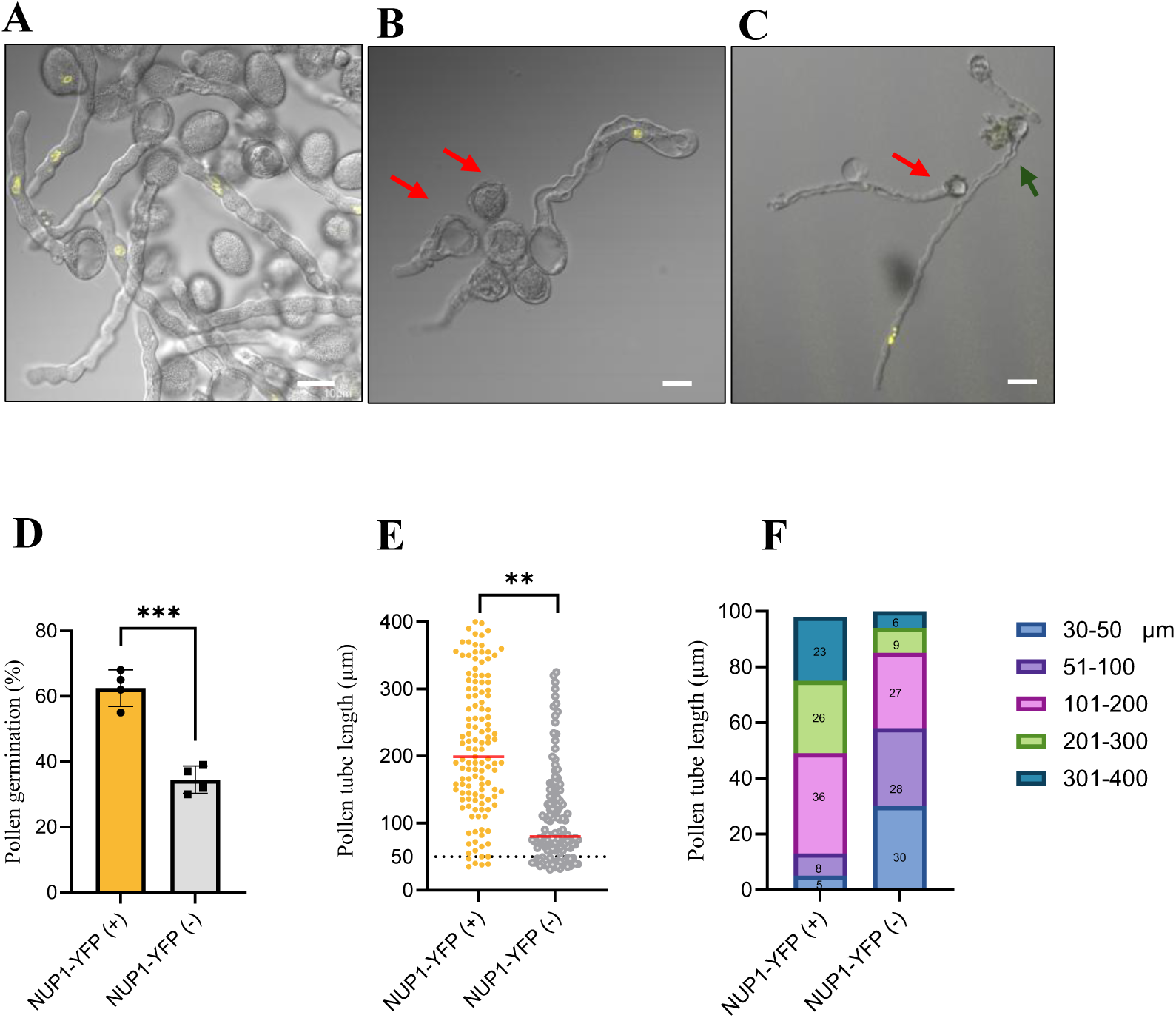
NUP1 is required for pollen germination and elongation of pollen tubes. A-C. A – The NUP1-YFP (+) and NUP1-YFP (-) pollen germination. B - A representative image showing lower pollen germination of NUP1-YFP (-) and C. A representative image showing shorter pollen tube length of NUP1-YFP (-). The red arrow indicates mutant pollen. Scale bar 20µm. **D.** Germination percentage of NUP1-YFP (+) and NUP1-YFP (-) pollen. N=200 **E.** Pollen tube length (PTL) of NUP1-YFP (+) and NUP1-YFP (-) pollen. N=100 **F.** Distribution of Pollen tube length (PTL) of NUP1-YFP (+) and NUP1-YFP (-) pollen. The PTL were divided into five categories. N=100 for each genotype. A two-tailed t-test was employed to test the significance of the difference in D and E. *<0.05, **< 0.01 and ***<0.001.

We also investigated pollens from *nup1-1* and *nup1-2^-/+^*plants, comparing them with those of Col-0 as a control (Supplementary Fig. 6). The PG of *nup1-1* was significantly lower (62 ± 4%) than that of Col-0 (71 ± 5%) and *nup1-2^-/+^* (67 ± 5%). The PTL of *nup1-1* (177 µm ± 22) and *nup1-2^-/+^* (192 µm ± 42) was significantly lower than that of Col-0 (205 µm ± 24). Altogether, NUP1 is required for pollen germination and pollen tube elongation.

### 4. NUP1 is highly expressed in pollen and maintains nuclear morphology

To examine NUP1 protein expression in pollen, we used the NUP1-YFP (+/+) transgenic plant, which produces pollen NUP1-YFP under the NUP1 native promoter. NUP1 protein is expressed in both VN and SN of all pollens, but its expression is much higher in VN than in SN (Figure 4A). Under low laser intensity, NUP-YFP is clearly visible in VN but barely seen in SN. However, under high laser intensity, it is visible in both. The fluorescence intensity profile shows that VN has a higher peak than SN (Figure 4B). Since NUP1 was previously reported to alter nuclear shape and size in Arabidopsis leaves and roots (Tamura & Hara-Nishimura, 2011), we hypothesize that it may also alter nuclear morphology in pollen. We compared the nuclear shape and size of pollen with and without NUP1-YFP expression by staining them with DAPI (Figure 4C). The NUP1-YFP pollen shows irregular/lobed VN, while the mutant pollen shows smaller and round VN. We further examined the VN shape in Col-0 and *nup1-1* pollen by DAPI staining (Figure 4D). The Col-0 has an elliptical VN, while *nup1-1* has a round one. Although the SN in the *nup1* mutant pollen and NUP1-YFP (-) pollen appear smaller and circular (Figure 4C-D), the SN was not studied because it was too small to accurately characterize the morphological change. Overall, NUP1 is required to maintain the nuclear shape and size in pollen specially the VN.

**Figure 4.**
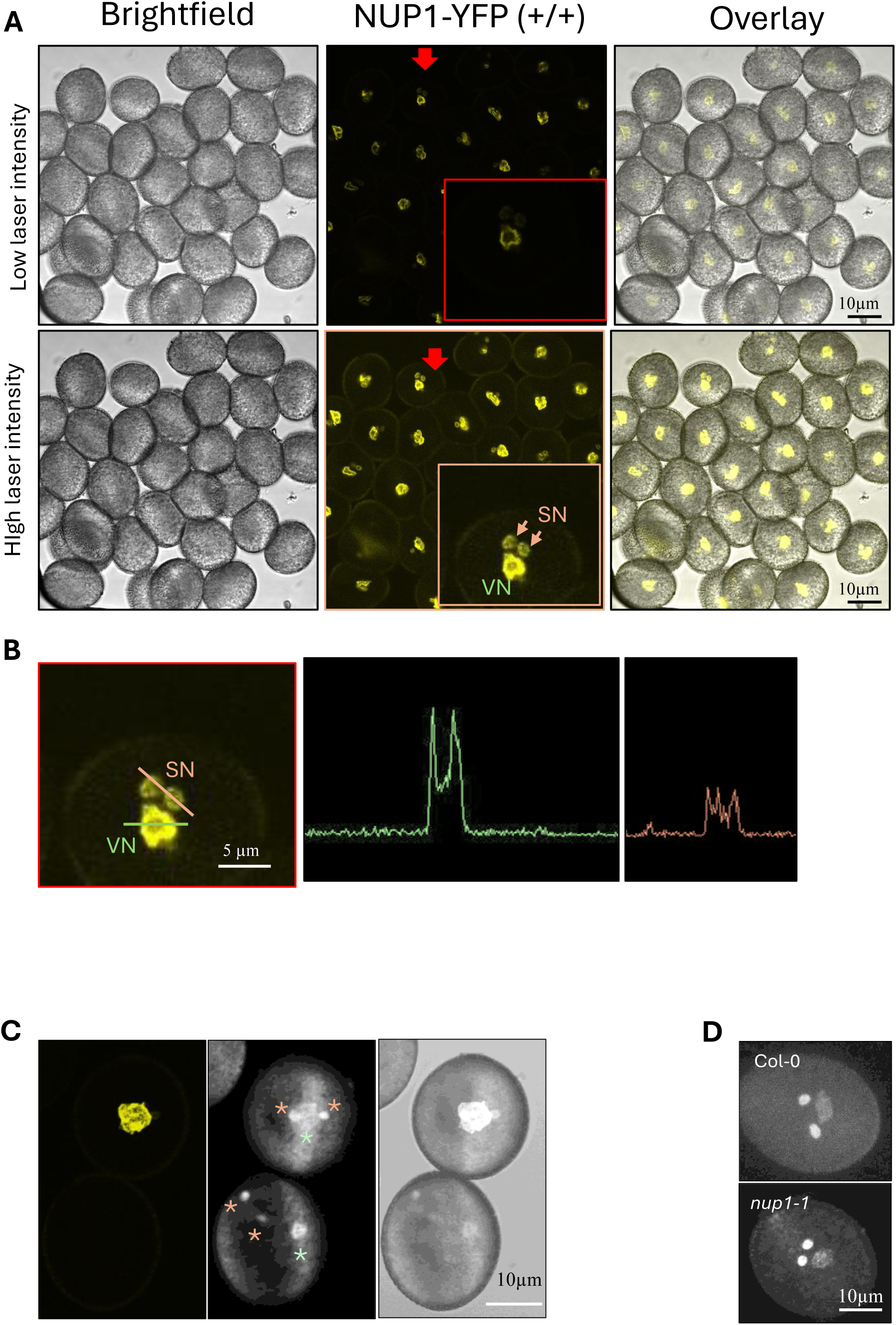
NUP1 is highly expressed in VN and maintains nuclear morphology. **A.** NUP1 expression in VN and SN. Image taken under low and high laser intensity. **B.** Relative fluorescence intensity of NUP1-YFP in VN and SN. **C.** Comparing the morphology of nuclei in NUP1-YFP (+) and NUP1-YFP(-) pollen. The green asterisk shows VN, and the orange asterisk shows SN. **D.** Comparing the nuclear morphology in Col-0 and *nup1-1* pollen.

### 5. NUP1 maintains the MGU organization in pollen

In addition to changes in VN shape and size in NUP1-YFP (-) pollen, we also noticed that it exhibits MGU organization that differs from that of NUP1-YFP (+) pollen (Figure 4C). To study the effect of a change in nuclear morphology on MGU organization, we examined pollen for nuclear arrangement (Figure 5A-F). Typically, MGU is organized with a VN at the center and SN on both sides, referred to as a connected/intact MGU as shown in NUP1-YFP (+) pollen (Figure 5A-B). The SN in NUP1-YFP (-) was far from the VN, and sometimes both SN were on the same side, referred to as disconnected MGU (Figure 5A-B). We also stained pollen from the *nup1-2*^-/+^ heterozygous plant with DAPI (Figure 5C-D). This plant produces ∼50% of the wild-type pollen and ∼50% of the NUP1-deficient pollen (*nup1-2*). We found two categories of MGU organization: one with a standard MGU organization/connected MGU (likely Wild type pollen), and the other with a disconnected MGU (likely *nup1-2* mutant pollen).

**Figure 5.**
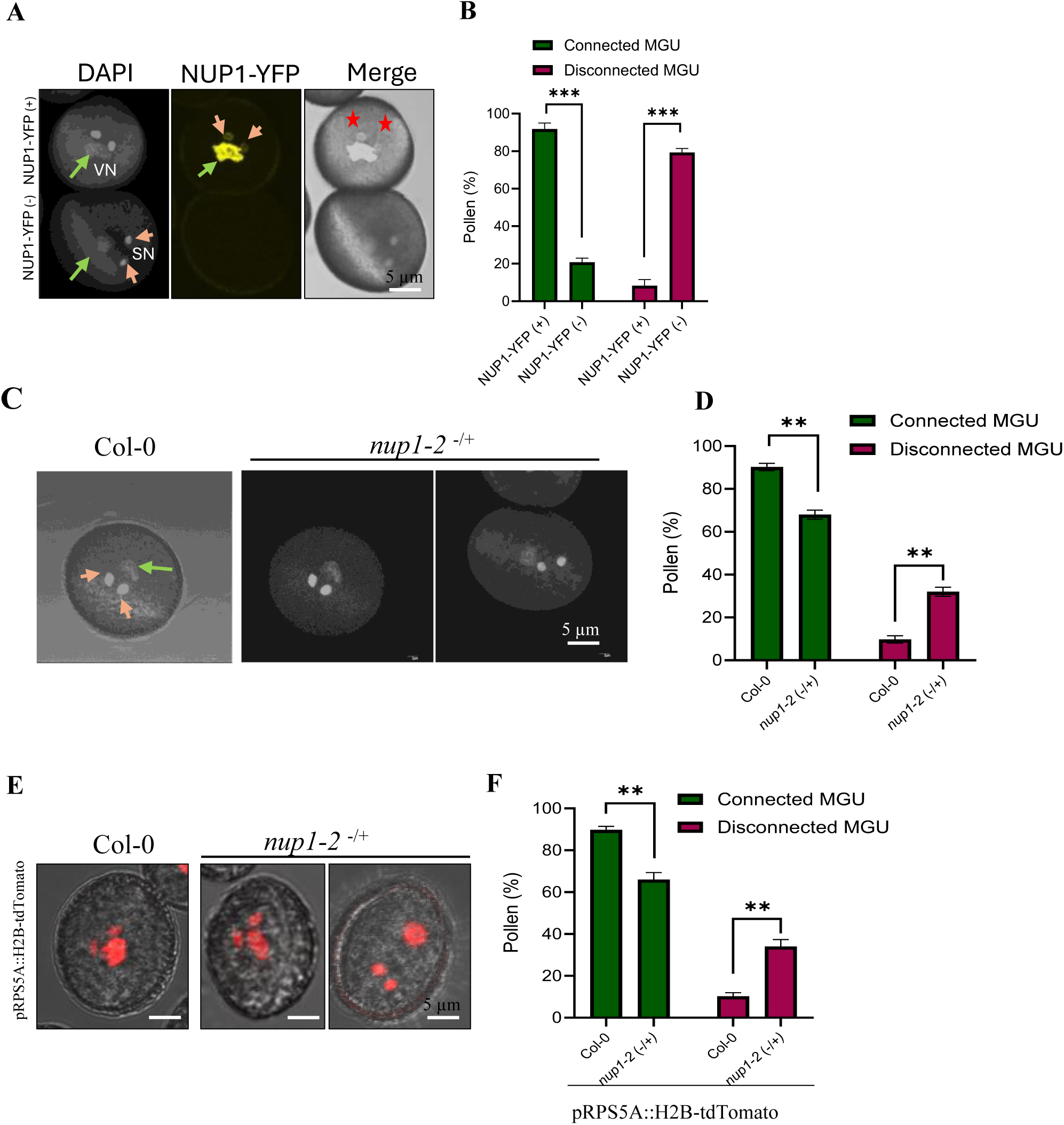
NUP1 is essential for MGU organization in pollen A-B. MGU organization in NUP1-YFP (+) and NUP1-YFP (-) pollen. N=50. Scale bar 5µm. **C-D.** MGU organization in pollen from *nup1-2* plant, which produces ∼50% wildtype and ∼50% *nup1-2* ko pollen, N=50, Scale bar 5µm. **E-F.** MGU organization in the transgenic line expressing pRPS5A::H2B-tdTomato in Col-0 and *nup1-2* background. N=100, Scale bar 5µm. A two-tailed t-test was employed to test the significance of the difference. *<0.05, **< 0.01 and ***<0.001. The green arrow shows VN, and the orange arrow shows SN.

Since NUP1 expression under its native promoter yields a weak fluorescence signal, especially in SN, we further generated new transgenic lines expressing *pRPS5A*:*H2B-tdTomato* (Maruyama et al., 2015) in the Col-0 and *nup1-2^-/+^* heterozygous plant backgrounds to study MGU (Figure 5 E-F). The *pRPS5A: H2B-tdTomato* Col-0 (hereafter referred to as H2B-Tom-Col) pollen has mostly standard MGU organization, which was similar to that observed in the Col-0 wild-type pollen. The *pRPS5A: H2B-tdTomato nup1-2*^-/+^ (hereafter referred to as H2B-Tom-*nup1-2^-/+^*) pollen had two categories of MGU organization: one standard, with intact MGU (∼60 %), and the other, with disrupted MGU (∼40%). Although the expected ratio was 1:1 for the MGU organization, the observed deviation may reflect stochastic inheritance of NUP1, which could partially compensate for the *NUP1* mutation in pollen. These results indicate that NUP1 is essential for maintaining MGU integrity.

### 6. The MGU migration is disrupted in *nup1* mutant pollen tubes

To determine whether changes in nuclear morphology of MGU also affect its transport, we conducted *in vitro* germination of NUP1-YFP (+/-) pollen and stained them with DAPI (Figure 6A). The MGU in NUP1-YFP (+) pollen was mostly near the pollen tube tip, as expected (N=17/20), while the MGU in NUP1-YFP (-) pollen was mostly disrupted with VN near the apex and SN in the pollen grain (N=13/16). Overall, the lack of NUP1 protein resulted in lower PG, shorter PTL and impaired MGU transport (Figure 6B). We could not obtain a higher number of pollen tubes suitable for assessing MGU position due to a dim fluorescent signal in NUP-YFP (+) pollen tubes. Also, the addition of DAPI diminishes the YFP signal, which could increase the risk of NUP-YFP (+) pollen being imaged as a false negative. Therefore, we conducted *in vitro* PG for the H2B-Tom-Col and H2B-Tom-*nup1-2*^-/+^ transgenic lines (Figure 7A-B and Supplementary Figure 7). The expected impaired MGU transport in H2B-Tom-*nup1-2*^-/+^ was ∼23% (Figure 7A), accounting for the fact that NUP1-deficient pollen has half the PG rate of that of Col-0, and ∼30% of the pollen tubes were too short to determine MGU position. In the H2B-Tom-Col-0 pollen tube, the MGU position was mostly near the apex of the pollen tube, and VN was leading the migration, followed by two SN (Figure 7B-D). These MGU positions and migration patterns are considered standard/normal based on past reports (Zhou & Meier, 2014). There was no change in MGU order in most of the H2B-Tom-*nup1-2*^-/+^ pollens (Figure 7C), but the MGU migration seems to be disrupted in almost 20% of the pollen tubes (Figure 7D). The ∼3% difference between observed and expected impaired MGU transport may be due to a large variation in PG and PTL. Based on observations of MGU in these two transgenic lines, we conclude that NUP1 is required for proper MGU transport in pollen tubes.

**Figure 6.**
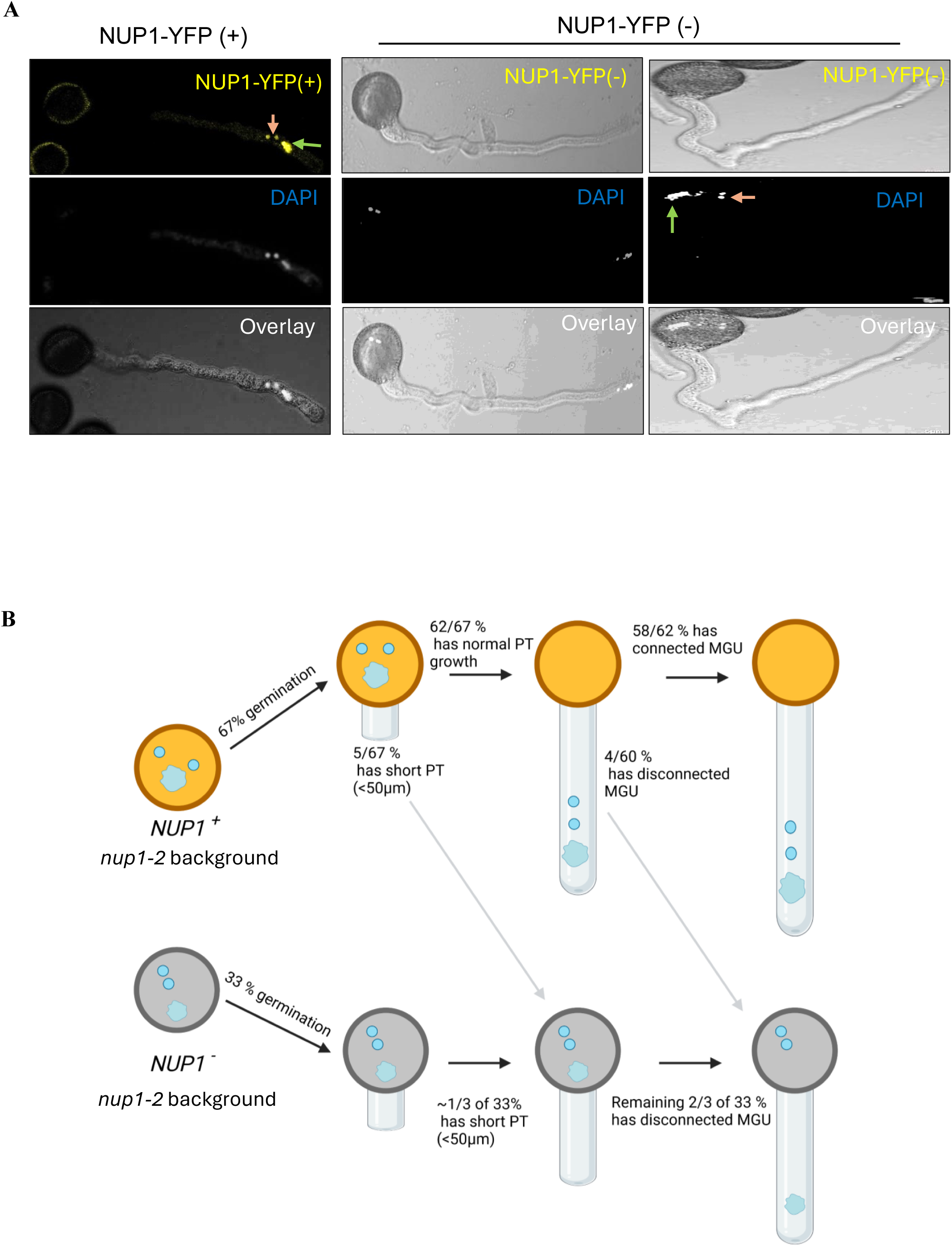
MGU positioning in NUP1-YFP (+) and NUP1-YFP (-) pollen tubes. **A.** MGU transport in NUP1-YFP (+) and NUP1-YFP (-) pollen tube. Intact MGU is at the pollen tube tip in NUP1-YFP (+), N=20, while VN is at the pollen tube tip and SN at the pollen grain in NUP-YFP (-), N=16. The orange arrow shows SN in the pollen, and the green arrow shows VN in the pollen tube tip of NUP-YFP (-). Scale bar 10µm. **B.** Effect of *nup1-2* mutation on pollen germination, pollen tube elongation and MGU. The *nup1-2* mutant pollen, lacking the *NUP1* gene, has a PG half compared to that of pollen with the *NUP1* gene. Therefore, out of 100 pollen, 67% of pollen has the *NUP1* gene, and 33% is the mutant. 1/3 of the germinated mutant pollen has a short PT (<50 µm), and the remaining 2/3^rd^ have an elongated PT but impaired MGU transport.

**Figure 7.**
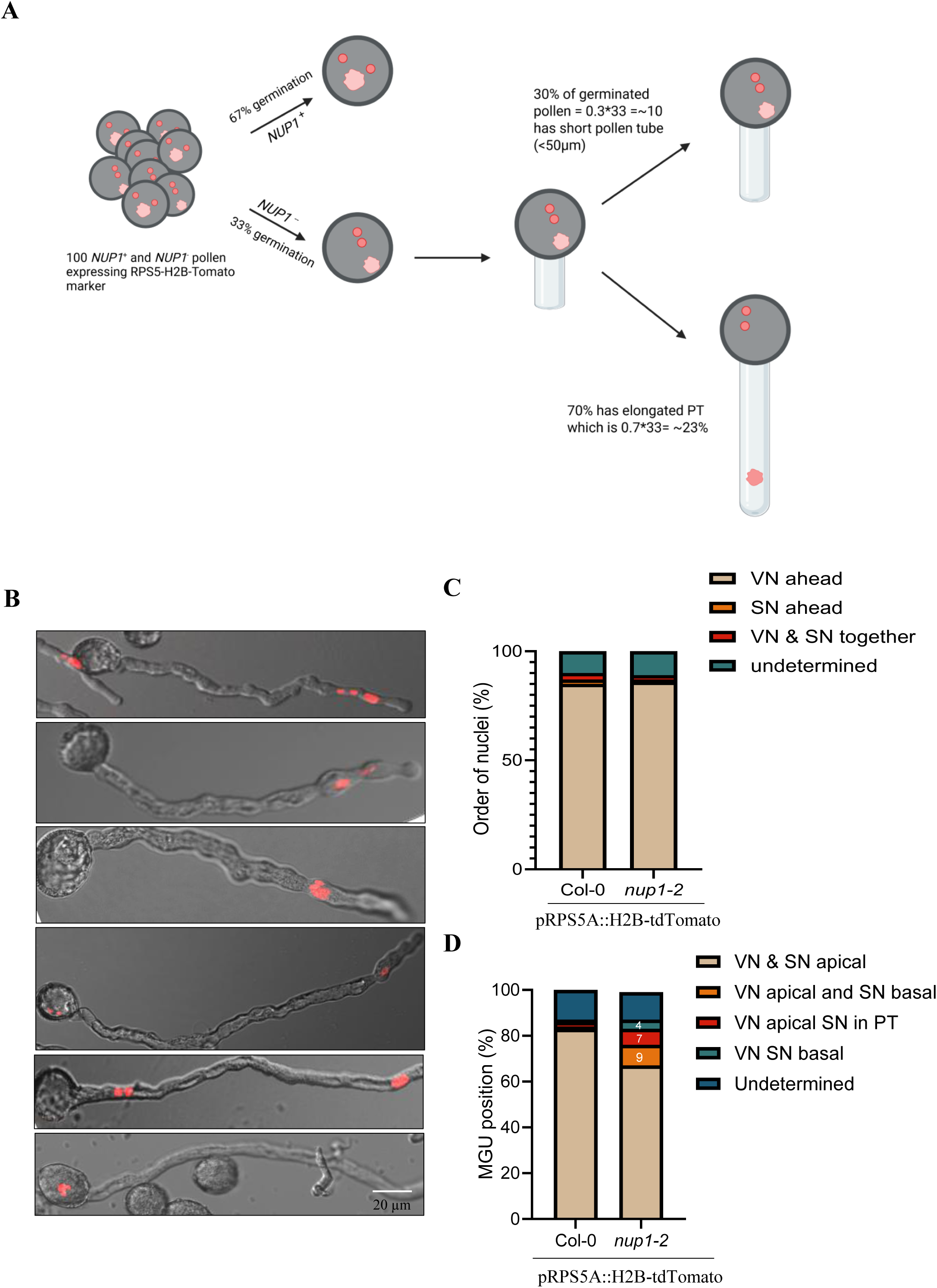
MGU positioning in pollen of Col-0 and *nup1-2^-/+^* plants expressing pollen nuclei marker. **A.** The MGU transport defect in H2B-Tom-*nup1-2*^-/+^ The *nup1-2*^-/+^ pollen has PG half than the pollen with *NUP1* expression. Therefore, out of 100 pollen, 67% of pollen have the NUP1 gene, and 33% are mutants (based on Fig 1j). Almost 30% of the mutant pollen has a short PTL (<50 µm) (Fig. 1k), which would be suboptimal for determining the MGU position. So, the remaining 70% of 33, which is ∼23, is the expected number of pollen with some form of MGU transport defect. **B-D.** The MGU transport in the transgenic line expressing pRPS5A::H2B-tdTomato in Col-0 and *nup1-2*^-/+^ background. B. Some representative images of MGU transport C. Quantitative analysis of the order of MGU in pollen tubes. D. Quantitative analysis of MGU position in pollen tubes. N=100, Scale bar 20µm.

## Discussion

The role of NUP1 in Arabidopsis male fertility has been reported by multiple groups, but the mechanistic details of how NUP1 loss affects pollen remain unknown. Here, we used the T-DNA lines *nup1-1* and *nup1-2^-/+^*and found that NUP1 is important for PG and PT elongation. The side-by-side comparison of pollens with or without NUP1 expression unexpectedly revealed that the NUP1 protein is inherited from the PMC to its daughter pollen cells. Interestingly, NUP1 was found to be highly expressed in VN and to regulate pollen nuclear morphology, especially the VN. More importantly, NUP1 was shown to be essential for the organization and transport of MGU in pollen tubes, likely through shaping the nuclear shape and size. Notably, our finding of nuclear morphology-mediated regulation of MGU may explain the mechanistic details underlying the defective MGU movement in previously reported mutants such as *kaku4*, *wit*, and *wip,* which have abnormal nuclear morphology (Goto et al., 2020; Zhou et al., 2015; Zhou & Meier, 2014).

We showed that NUP1 is required for male fertility by maintaining pollen morphology and PG, which is in agreement with previous studies (Bao et al., 2019; Tamura et al., 2010). The PG and pollen tube elongation would ideally require a large amount of mRNA to make abundant proteins to support the rapid growth, but the mutation in *NUP1* would slow down the mRNA export (Lu et al., 2010). This might be the reason the mutant has lower PG and PTL than the wild type. Future work could focus on isolating and analyzing the nuclear-trapped mRNA in mutant pollen.

There was a discrepancy between the expected genotype of the pollen tetrad described in Figure 2A and the results presented in Figure 2B. This suggests that, even though half of the daughter pollen cells in the tetrad were genetically mutant (lacking a functional copy of NUP1-YFP), they still inherited some NUP1 proteins from their PMCs. Notably, two of the pollen had brighter fluorescence, and the other two had a light-yellow colour, indicating the protein dilution during inheritance. NUP1 appears to be continuously diluted during maturation; ultimately, in mature pollen grains, YFP fluorescence was completely lost in half of them. The NUP1 protein produced in PMC is inherited during meiosis and likely compensates for a genetic NUP1 deficiency in mutant pollen, at least in maintaining pollen morphology. Once the inherited NUP1 protein is diluted and pollen begins to germinate, the effects of the *NUP1* mutation gradually unfold. The mutant pollen has lower PG, shorter PTL, disconnected MGU, and impaired MGU transport compared to the wild type. Together, these series of defective events completely block the *nup1-2* pollen tube from delivering sperm cells to the ovule, which explains why we never obtained *nup1-2* homozygous seeds.

The MGU’s disconnection and impaired transport in the mutant pollen tube may result from altered nuclear morphology. NUP1 was previously reported to regulate the nuclear morphology in Arabidopsis leaves and roots (Tamura et al., 2010), and MGU has been shown to be connected by microtubules (Åström et al., 1995; Wang et al., 2024). Several studies suggest that microtubules physically link VN and SC and are required for proper MGU migration from the pollen grain to the apex (Wang et al., 2024). Therefore, it would be interesting to examine potential interactions between NUP1 and microtubules and their effects on nuclear morphology and MGU. We speculate that the NUP1 protein in the nuclear envelope may serve as an anchor for microtubules that connect VN and SN.

MGU migrates from the pollen grain to the tip of the pollen tube, but the mechanism driving this journey remains poorly understood. One conjecture is that cytoplasmic streaming moves all organelles toward the tip (Chebli et al., 2013) and is driven by the motor protein myosin (Buchnik et al., 2015). Recently, some proteins have been implicated in controlling the movement of VN and SN in pollen tubes. KAKU4, a putative nuclear laminar protein, was shown to regulate the nuclear shape in vegetative cells. KAKU4 also ensures the orderly migration of the MGU in the elongating pollen tubes (Goto et al., 2020). Similarly, two families of nuclear membrane proteins, WPP domain-interacting proteins (WIPs) and WPP domain-interacting tail-anchored proteins (WITs), are required for maintaining the correct order of the nuclei in the pollen tube (Zhou and Meier, 2014). Interestingly, proteins involved in MGU movement, such as KAKU4, WIPs, and WITs, also regulate nuclear shape and size (Goto et al., 2020), suggesting an intricate relationship between nuclear morphology and the orderly movement of the MGU. In conclusion, we found that NUP1 is important for several aspects of pollen development, particularly in MGU organization and transport, likely through regulation of VN morphology. Understanding the mechanistic details of MGU function may be useful for developing heat-resistant pollens in the future amid rising heat waves.

## Materials and Methods

### Plant material and growth conditions

Arabidopsis wild type (Col-0) and the T-DNA mutant *nup1-2^-/+^*(SALK_020221) were all obtained from Arabidopsis Biological Resource Center (ABRC). The genotyping primers used to characterize the mutant lines are listed in Supplementary Table 4. Plants were grown in a walk-in growth room at 23 °C, 150 μmol/m^2^/s light intensity, and 50% humidity under 16-hr light/8-hr dark conditions.

### Plasmid construction and transgenic plants

A transgenic line expressing NUP1 fused with YFP driven by its native promoter in *nup1-2* background (*nup1-2 pNUP1::NUP1-YFP*) was previously generated (Lu et al., 2010; Thapa et al., 2022). The pRPS5A:H2B-tdTomato plasmid (Maruyama et al., 2015) was transferred to *Agrobacterium tumefaciens* GV3101 through electroporation. Then, the Agrobacterium-mediated floral dip method was used to generate transgenic lines in the Col-0 and *nup1-2* backgrounds, which were subsequently verified by kanamycin selection, PCR genotyping, and fluorescence microscopy.

### DAPI (4′,6-Diamidino-2-phenylindole dihydrochloride) staining of pollen

Pollen nuclei were stained with DAPI (Sigma #D9542, Mississauga, Canada). 1µg/mL of DAPI solution was prepared in pollen isolation buffer (PIB) containing 100 mM NaPO_4_, pH 7.5, 1 mM EDTA, and 0.1 % (v/v) Triton X-100. The healthy pollen was isolated from 4-6-week-old *Arabidopsis* plants and suspended in 1 mL of PIB. The pollen-containing buffer was briefly centrifuged at 1,500 g to pellet the pollen. Finally, the pollen was resuspended in 50 µL PIB and incubated at room temperature for 10 min. Pollen nuclei were viewed with a fluorescence microscope using the DAPI filter set.

### Confocal microscopy and imaging

Imaging was performed using a confocal microscope (Olympus FV1000). The following conditions were used to detect various signals in pollen and pollen tubes. For YFP detection, excitation 515 nm, emission 520-550 nm; for DAPI, excitation 405 nm, emission 470-500 nm; for tdTomato, excitation 554 nm, emission 575-585 nm.

### *In vitro* pollen germination assay

To measure pollen germination (PG) efficiency, pollen from a healthy plant (4-6 weeks old) was obtained by isolating anthers. Pollen was germinated on *in vitro* pollen medium containing 5 mM 2-(N-morpholino) ethanesulfonic acid (MES) (pH 5.8 adjusted with Tris), 1 mM KCl, 10 mM CaCl_2_, 0.8 mM MgSO_4_, 1.5 mM boric acid, 16.6% (w/v) sucrose, 3.65% (w/v) sorbitol, and 1% (w/v) agar (Fan et al., 2001). Pollen medium (200-300 µL) was gently poured onto clean microscope slides and left at room temperature for 5 min to solidify. Anthers were gently brushed on medium to spread the pollen. The slides were incubated at 20-25°C for 8 hrs in the dark, at room temperature and high humidity. After that, PG was observed under a microscope.

### Statistical analysis

Means, Standard Deviations (SD), and Standard Error of Mean (SEM) were calculated using GraphPad Prism 10. All experiments were performed using at least three biological replicates. Statistical analysis was performed by Student’s t-test. A p-value of 0.05 or less was used as a statistically significant difference indicated by *, a p-value of 0.01 or less was denoted by **, and a p-value of 0.001 or less was denoted by ***. All the graphs were generated using GraphPad Prism 10.

## Accession number

The gene ID of NUP1/NUP136 is At3g10650.

## Acknowledgments

We thank the Arabidopsis Biological Resource Centre (ABRC) for providing the mutant seed used in this study. We thank Daisuke Maruyama and Tetsuya Higashiyama for providing the *pRPS5A::H2B-tdTomato* plasmid. This work was supported by grants from the Natural Science and Engineering Research Council of Canada (RGPIN/04625-2017, to Y.C.); and Agriculture and Agri-Food Canada (to Y.C.). X.X was supported by a scholarship from the China Scholarship Council.

## Author contributions

YC, GT and RKT conceived the project. YC, RKT, GT, and SEK designed the experiments. RKT and GT crossed the NUP1-YFP (+/+) with *nup1-2* ^-/+^. BS and XX generated the transgenic line expressing *pRPS5A:H2B-tdTomato in the Col-0 and nup1-2^-/+^ backgrounds*. JS, CC and SB conducted the PCR-based genotyping. BS and RKT performed the PG assay. All other experiments were conducted by RKT. XL, SD, RKT and SJR analyzed the data. RKT wrote the first draft, and YC revised and edited the manuscript. All authors read and approved the final article.

## Supplementary Tables

**Supplementary Table 1:** Cross using wild-type as pollen donor and *nup1-2^-/+^* as pollen recipient.

**Supplementary Table 2:** Reciprocal cross using *nup1-2^-/+^* as pollen donor and wild-type as pollen recipient.

**Supplementary Table 3:** Segregation in *nup1-2^-/+^* heterozygous selfed F2 seeds.

**Supplementary Table 4:** List of primers used in this study.

