## Supplementary Figures 1-7 for "Nucleoporin1 maintains male germ unit organization and transport in Arabidopsis pollen tubes, likely through shaping nuclear morphology"

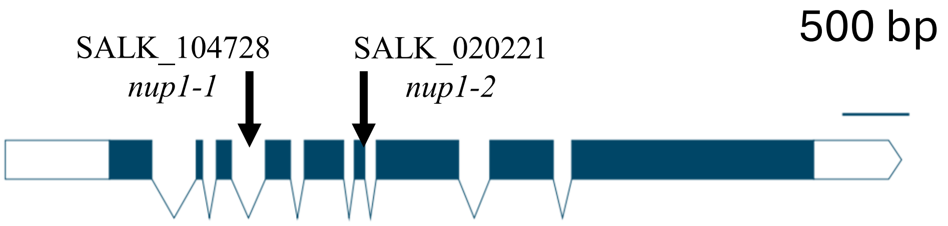


**Supplementary Figure 1. The *NUP1* gene map showing the intron (line break) and exon (blue box).**
The sites of T-DNA insertion for two mutants, *nup1-1* and *nup1-2*, are shown by an arrow.

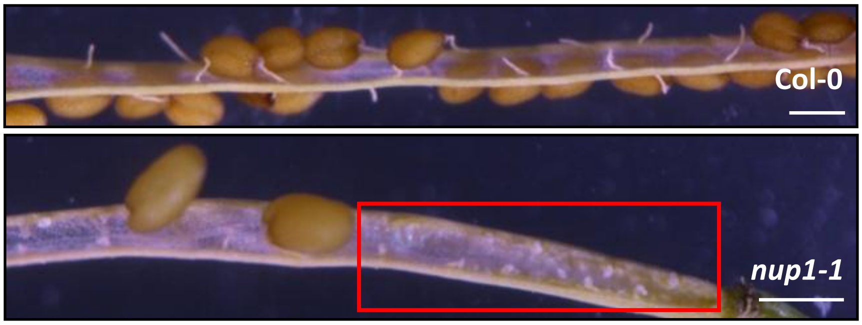


**Supplementary Figure 2. Mature siliques of Col-0 and *nup1-1*.**
The silique from mutant plants has many unfertilized ovules (red box), which are absent in Col-0.

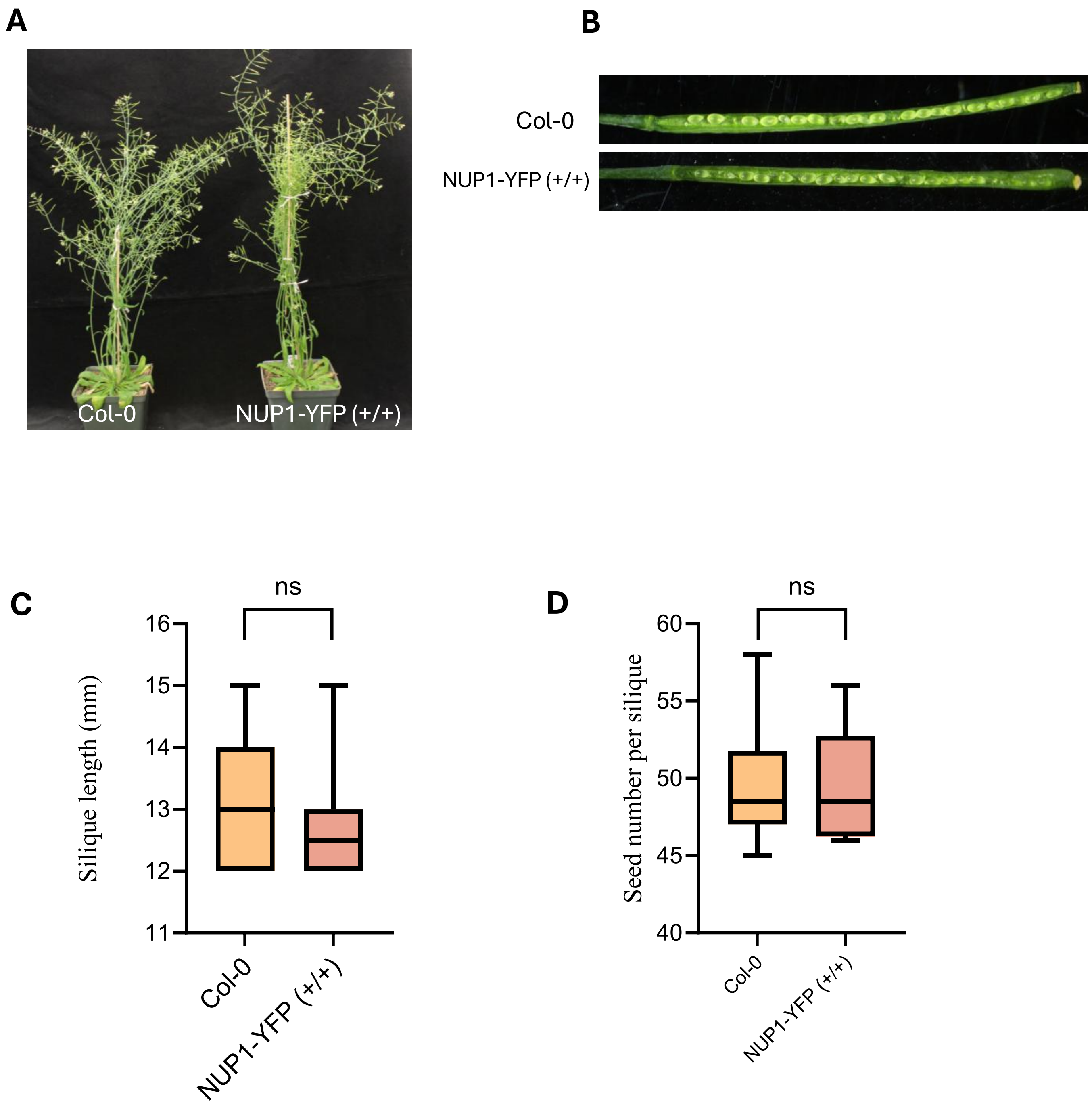


**Supplementary Figure 3. Complementation of *nup1-2*$^{-/+}$ Arabidopsis plant with NUP1-YFP transgene**

**A.** Mature plant Col-0 and NUP1-YFP (+/+)

**B.** Comparison of silique from Col-0 and NUP1-YFP (+/+) plant. N=20 for each genotype.

**C-D.** Comparison of silique length and seed number in Col and NUP1-YFP (+/+) plants. N=20 for each genotype. A two-tailed t-test was employed to test the significance of the difference. ns = no significant difference, *<0.05, **< 0.01 and ***<0.001.

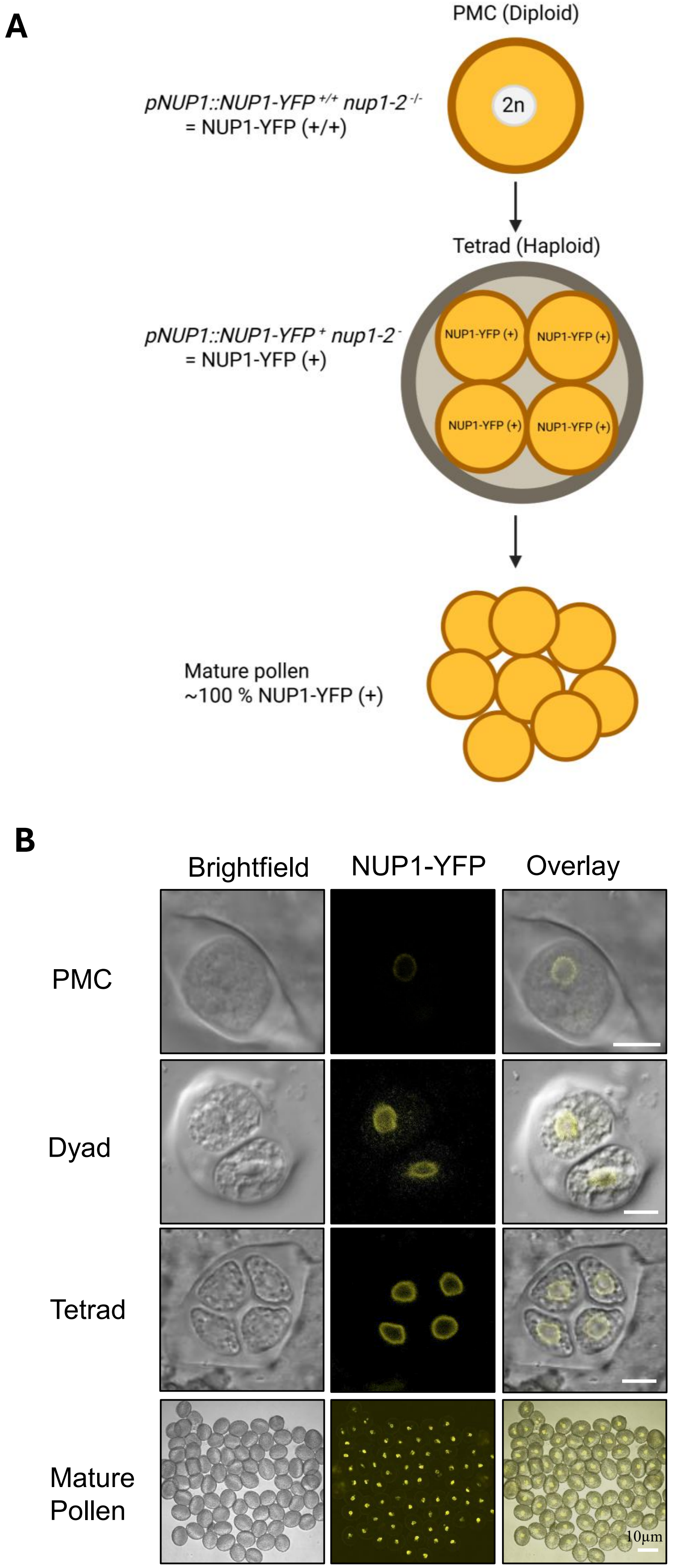


**Supplementary Figure 4. Arabidopsis male gametogenesis in NUP-YFP (+/+) plant.**
**A.** Schematic diagram of male gametogenesis showing PMC to mature pollen
**B.** NUP1 expression in PMC to the mature pollen stage
Scale bar 10µm.

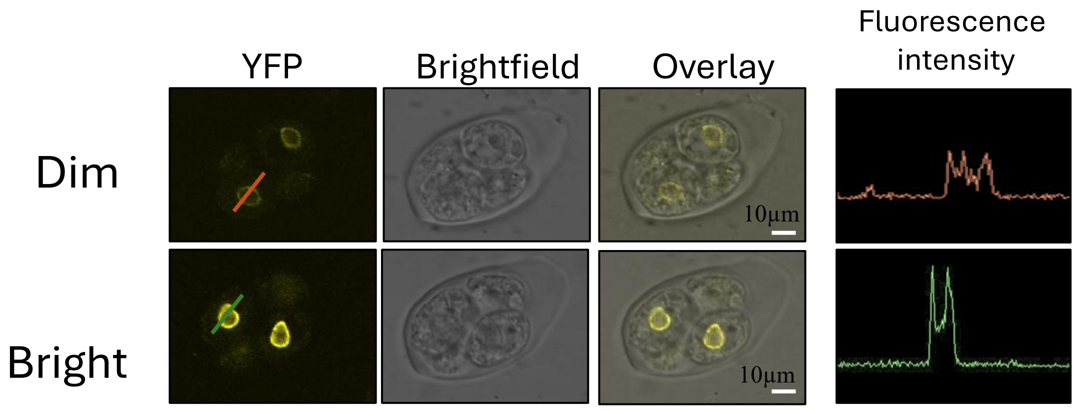


**Supplementary Figure 5. NUP1-YFP expression in Arabidopsis pollen (NUP1-YFP (+/-) tetrad.**

NUP1-YFP is expressed in all four cells of the tetrad, although it was expected to be seen in only two of them. However, it is highly expressed (bright) in two of them and lowly expressed (dim) in the remaining two, indicating inheritance of the NUP1 protein during gametogenesis. Both dim and bright images were taken under identical conditions and with the same laser intensity. Scale bar 10μm.

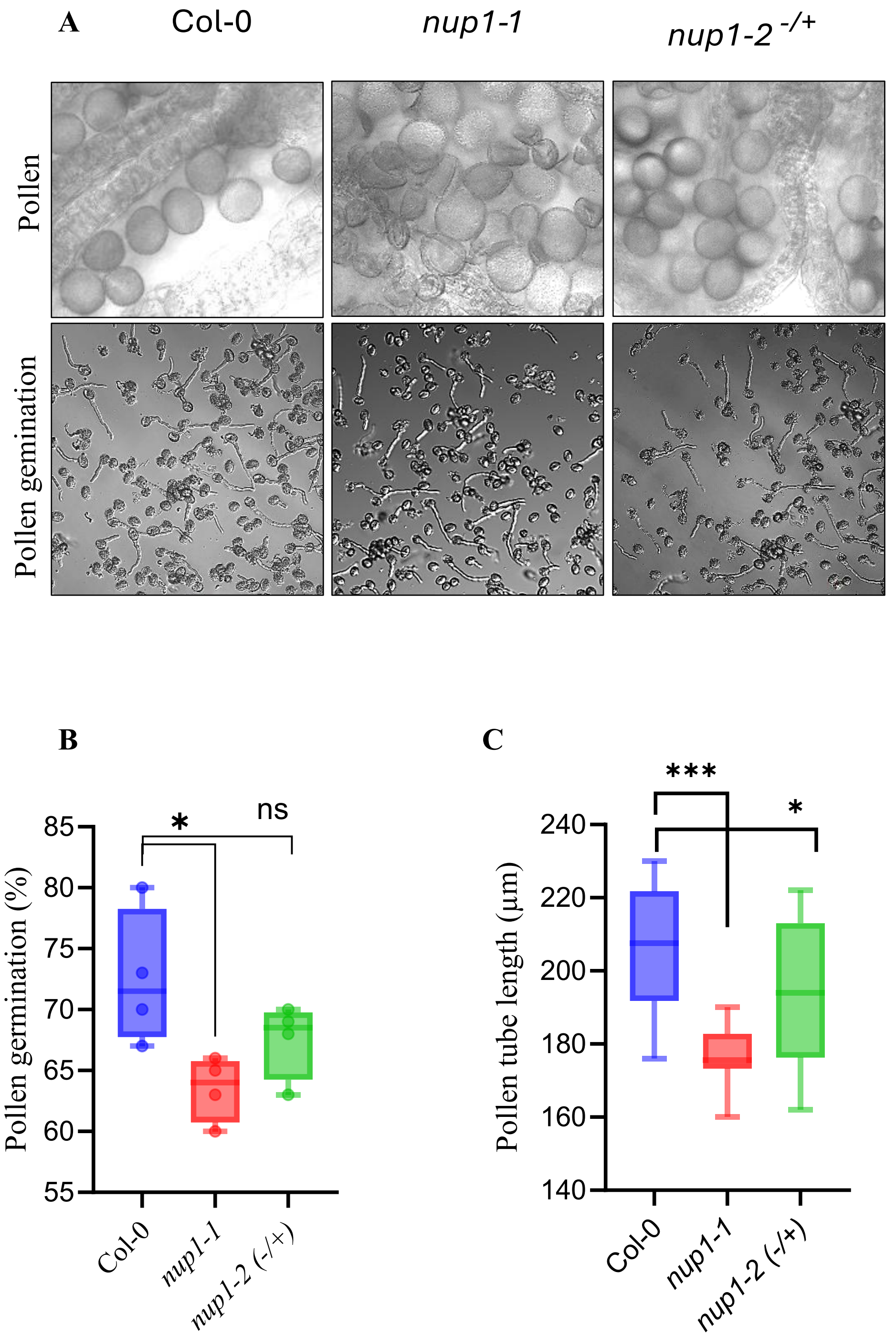


**Supplementary Figure 6. Comparing pollen germination and pollen tube length in *nup1* mutants**

**A.** Comparing pollen of Col-0, *nup1-1* and *nup1-2* $^{-/+}$ plants.

**B.** Pollen germination percentage in different genotypes.

**C.** Pollen tube length in different genotypes.

N = At least 100 for each genotype and at least three biological replicates for each experiment.

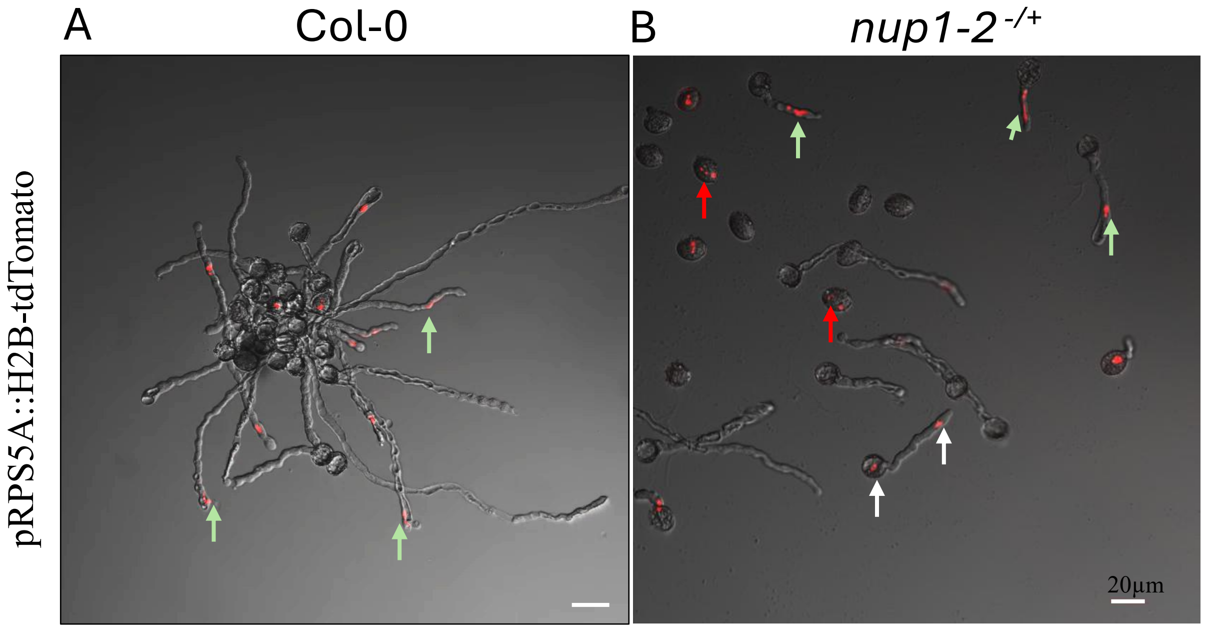


**Supplementary Figure 7. Pollen germination from RPS5-H2B-Tom-Col-0 and RPS5-H2B-Tom-*nup1-2* $^{-/+}$ plant.**

The green arrow shows an intact MGU at the pollen tube tip, the red arrow shows a dissociated MGU in a pollen grain, and the white arrow shows a disconnected SN and VN. Scale bar 20µm. N=100 for each genotype.
